# Protein-surfactant-polysaccharide nanoparticles increase the catalytic activity of an engineered β-lactamase maltose-activated switch enzyme

**DOI:** 10.1101/746560

**Authors:** J.P. Fuenzalida, T. Xiong, B. M. Moerschbacher, Marc Ostermeier, F.M. Goycoolea

**Author notes:** **Corresponding Authors F.M. Goycoolea**, (F.M.G.), **Marc Ostermeier** (M.O.).

## Abstract

We present polysaccharide-based nanoparticles able to associate and increase the catalytic activity of the maltose-binding MBP317-347 switch enzyme. Fluorescence quenching and molecular docking studies along with the partial resistance to increasing pH and ionic strength indicate that the increase in enzymatic activity is due to a specific interaction between the maltose binding pocket on MBP317-347 and alginate exposed on the surface of the nanoparticles. Finally, we show that the hybrid self co-assembled particles increase the half-life of MBP317-347 over six-fold at 37°C, thus reflecting their potential use as a macromolecular drug delivery system.

## Introduction

Enzyme delivery using nano-sized carriers is a promising research field since recent studies on nanotechnology led to the synthesis of uniformly sized, and non-cytotoxic nanoparticles1. Among the repertoire of available nanomaterials, soft nanomaterials based on protein and polysaccharides represent an enormous opportunity since several of them are biocompatible, biodegradable and generally regarded as safe (GRAS) according to the FDA.^1, 2^

Enzymes and proteins can be associated/immobilized either onto the surface or within the matrix of polymeric nanoparticles.^2-5^ Either of these techniques improve the stability of the cargo, as they serve to protect it against degradation. However, it has also been reported that enzyme immobilization in nanoparticles also often reduces enzyme activity.^6^

In a preceding paper, we described the preparation and physicochemical properties of an alginate-lysozyme nanoparticles capable of associating electrostatically with β-lactamase.^7, 8^ Here we describe the development of an alginate-based nanoparticle combined with lysozyme, albumin, and lecithin to stabilize the MBP317-347 switch enzyme. MBP317-347 is an engineered allosteric enzyme comprising a fusion of the maltose binding protein (MBP) and TEM1 β-lactamase (BLA) to create a β-lactamase enzyme whose catalytic activity is modulated by the presence of maltose9. Analogous switch enzymes might be used to activate a pro-drug in the presence of a specific effector, e.g. a high concentration of lactate in tumors.^10, 11^ In soft nanomaterials the interaction between enzymes and its constituents normally is restricted to non-specific interaction forces such as electrostatic, hydropohibic and van der Waals.

In an accompanying paper we have addressed the preparation and stability of nanoparticles comprised by alginate, lysozyme, albumin and lecithin (ref). In the present study, we exploit the similarity between the structure of maltose and alginate to facilitate the interaction between the nanoparticle and MBP317-347.12 We evaluated the properties of the maltose binding domain of MBP317-347 switch enzyme in the overall interaction between the switch enzyme and the alginate present in the different nanoparticles of varying composition. We find that nanoparticle association improves the catalytic activity and stability of MBP317-347. We provide proof-of-concept of the application of this system as a functional protein nanocarrier for the potential delivery of therapeutic enzymes.

## Materials and methods

### Materials

Seaweed sodium alginate samples were supplied by Danisco© (Denmark). The M/G ratio was 1:11 and the MW was ∼198 kDa (according to the manufacturer’s specifications). Lysozyme, human albumin, and other reagents of high purity were purchased from Sigma-Aldrich (München, Germany).

## Methods

### Preparation of the nanoparticles

The general scheme for the preparation of three different systems is shown in Figure 1. Nanoparticles comprising alginate and lysozyme were prepared at room temperature by mixing defined volumes of each component in solution. Briefly, 1 mL 23.3 mM alginate was added to 14 mL 0.043 mM lysozyme with gentle magnetic stirring (pH 4.5, 40 mM NaCl). The final concentrations of alginate and lysozyme were 1.56 and 0.041 mM, respectively. At pH 4.5, lysozyme carries 11 positive charges, 13 so the equivalent positive charge concentration (n^+^) was estimated to be 0.45 mM and the corresponding negative charge concentration of alginate (n^−^) was estimated to be 1.56 mM. The total charge ratio (n+/n^−^) was therefore always 0.29 with fixed total amount of charge of 2 mM.

**Figure 1.**
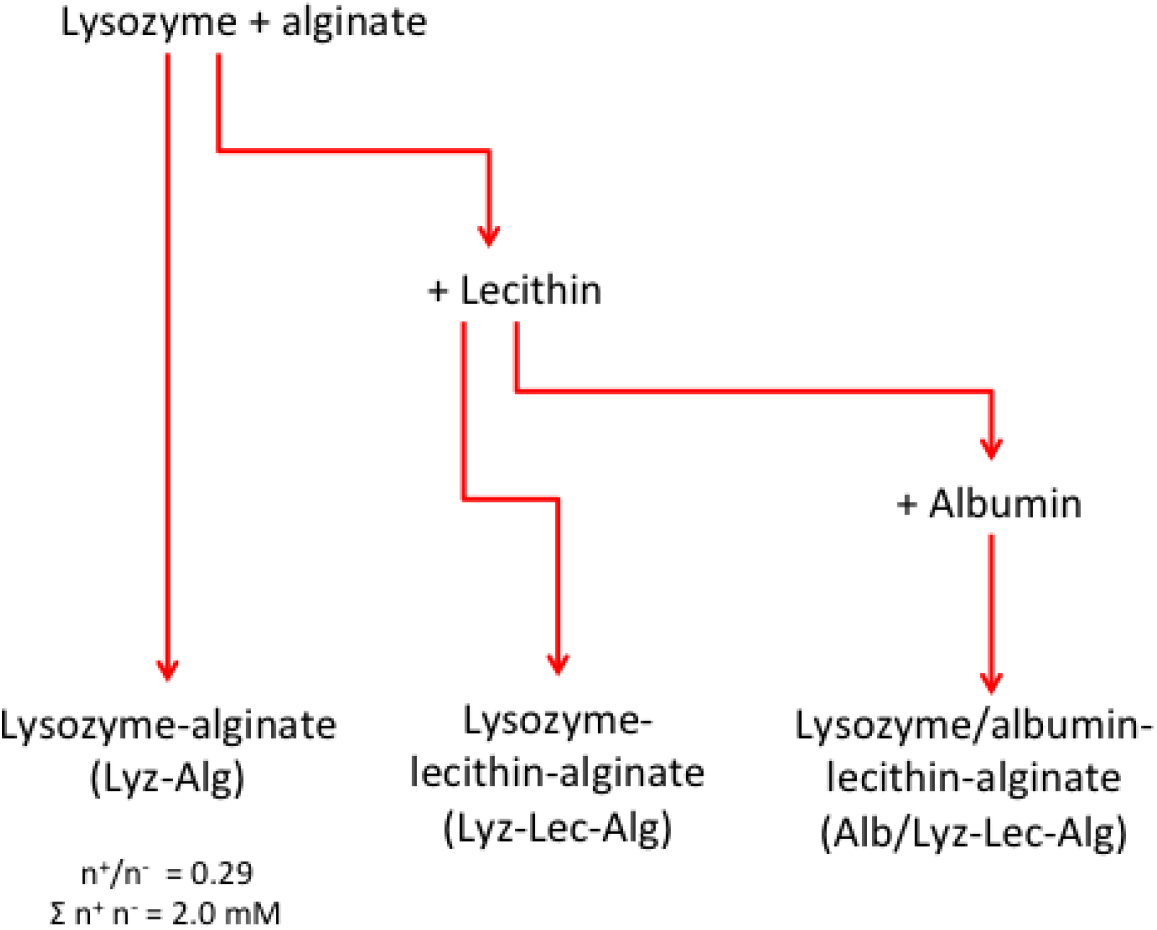
Overall scheme for the preparation protocol of the lysozyme/alginate-based nanosystems

Nanoparticles comprising alginate, lysozyme and lecithin were prepared by adding 220 μL 120 mg/ml lecithin in ethanol to lysozyme, with gentle stirring, prior to the addition of alginate as described above. The concentration of added lecithin was calculated to achieve ∼50 lipid molecules per lysozyme molecule.

Nanoparticles comprising alginate, lysozyme, lecithin and albumin were prepared by the addition of albumin to lysozyme before the addition of lecithin as described above, resulting in the replacement of 10% of the lysozyme charges with albumin charges. Finally, alginate was added as described above. At pH 4.5, albumin carries 16 positive charges.^14^ The final concentrations of lysozyme and albumin were 0.036 and 0.027 mM, respectively.

### Physical characterization

The particle size distribution was determined using two methods: dynamic light scattering with noninvasive back scattering (DLS-NIBS) and nano-tracking analysis (NTA). DLS-NIBS was carried out using a Malvern Zetasizer NanoZS (ZEN 3600, Malvern Instruments, Worcestershire, UK) fitted with a red laser (λ = 632.8 nm). NTA was carried out using a NanoSightTM LM10 system equipped with a LM14 green (535 nm) laser module and a cooled Andor camera (Andor-DL-658-OEM). The particles were diluted 1:100000 in water before analysis. The value of the surface zeta potential (ζ) was determined by mixed laser Doppler electrophoresis and phase analysis light scattering (M3-PALS) in milli-Q water and also using the ZetaSizer NanoZS instrument.

### MBP317-347 production

The design, synthesis and characterization of the MBP-BLA switch enzyme MBP317-347 were described previously.^9^ Briefly, *Escherichia coli* strain BL21(NEB) cells were transformed with the corresponding vector and grown overnight at 37°C in medium containing 6.7 mg/mL Na_2_HPO_4_, 3 mg/mL KH_2_PO_4_, 0.5 mg/mL NaCl, 1 mg/mL NH_4_Cl, 2 mM MgCl_2_, 0.1 mM CaCl_2_, 0. 03 mM thiamine-HCl, 50 mM fructose, 0.2% glycerol and 0.05 mM chloramphenicol. Protein expression was induced overnight at 20°C with 1 mM IPTG and the cells were lysed allowing the purification of MBP317-347 by affinity chromatography using a His-Tag column (GE Healthcare Life Science). The MBP317-347 protein was eluted with imidazole, dialyzed at 4°C and concentrated to ∼6 mg/mL in PBS (pH 7.4) containing 20% glycerol.

### Measurement of MBP317-347 activity

The enzymatic activity of MBP317-347 was determined as previously described using a nitrocefin hydrolysis assay.^15^ Maltose-induced switching was measured at 25°C in the presence of different effectors (maltose, alginate, and different nanoparticle formulations). The initial rate of nitrocefin hydrolysis was monitored at λ = 486 nm as previously described.^16^ Enzyme activity was measured in 30 mM phosphate buffer at pH 5.0 or 7.0 and in the presence of 12 or 100 mM NaCl. The effectors were added at the same total disaccharide concentration of 233 μM, and the concentrations of protein and nitrocefin were 50.9 nM and 50 μM, respectively. The samples were equilibrated for 20 min at 30°C before adding nitrocefin. Enzymatic rates were also measured as a function of effector concentrations and the data were fitted to a one-site-specific binding model using GraphPad Prism v6.0 to determine the affinity constants. The impact of 39 μM maltose on MBP317-347 activity in the presence of different nanoparticles was measured using 8.4 nM MBP317-347 and 8 μM nitrocefin in 30 mM phosphate buffer (pH 5.0) containing 12 mM NaCl, with each formulation presented at a disaccharide concentration of 39 μM.

### Quantitative SDS-PAGE

The retention of MBP317-347 by the Lyz-Lec-Alg nanoparticles was determined indirectly. Briefly, the above-described solution of MBP317-347 in presence of Lyz-Lec-Alg nanoparticles were centrifuged for 45 min at 10 000 × g using a Mikro 220R centrifuge (Andreas Hettich GmbH, Tüttlingen, Germany). The supernatants were carefully separated and loaded against a calibration curve of MBP317-347 in a denaturing SDS page (NuPAGE^®^ Tris-Acetate Pre-Cast gels). Loading and staining of the sample were performed according to the manufacturer’s recommendations. (Life Technologies, Grand Island, NY, USA; Protocol number MAN0007896 Rev 1.0). Precision Plus Protein™ All Blue (Bio-Rad Laboratories, Hercules, CA) was used as molecular weight sizing standard. Analytical analysis of the protein concentration in each sample was perform using ImageJ (National Institutes of Health) software analysis, as describe by Carter et al.^17^

### Thermal stability

The thermal stability of MBP317-347 was evaluated at 37° and 4°C in 50% PBS containing, 10% glycerol and 50.9 nM MBP317-347, with or without 233 μM maltose and with the nanoparticles present at a disaccharide concentration of 233 μM. Independent 20-μL aliquots were taken at different time intervals and the enzymatic activities were measured in 100 μL 100 mM phosphate buffer (pH 7.0) containing 233 μM carbohydrate ligand

### Molecular modeling

Molecular docking calculations for the association between maltose and MBP in the different nanoparticle formulation were prepared using Autodock v4.2. The crystal structure of MBP as a complex with maltose was retrieved from the RCSB Protein Data Bank (PDB entry 1ANF).

## Results and Discussion

MBP317-347 is an engineered switch protein with the maltose-binding protein (MBP) as the input domain and TEM1 β-lactamase (BLA) as the output domain. Maltose induces a conformational change that activates BLA.^9, 16^ Maltose comprises two glucose units joined by an α(1-4) glycosidic bond whereas alginate is a co-polymer that comprises M-blocks of β-(1-4)-linked poly-D-mannuronate, G-blocks of α(1-4)-linked poly-L-guluronate and MG-blocks of alternating residues. As a first test to test if the interaction between alginate and the maltose binding domain is possible, a tryptophan quenching experiment was performed. The fluorescence quenching effect of alginate (Figure 2) strongly suggests the existence of an interaction between alginate and MPB317-347. This kind of signal quenching normally reflects conformational changes in the protein induced by the presence of specific ligands, interaction of the ligand with the tyrptophans, or protein denaturation.^18^ The difference in the magnitude between the quenching caused by maltose and alginate suggests a strong conformational change in presence of the polysaccharide that is distinct from that induced by maltose and confirmed a binding interaction between alginate and MBP317-347.

**Figure 2.**
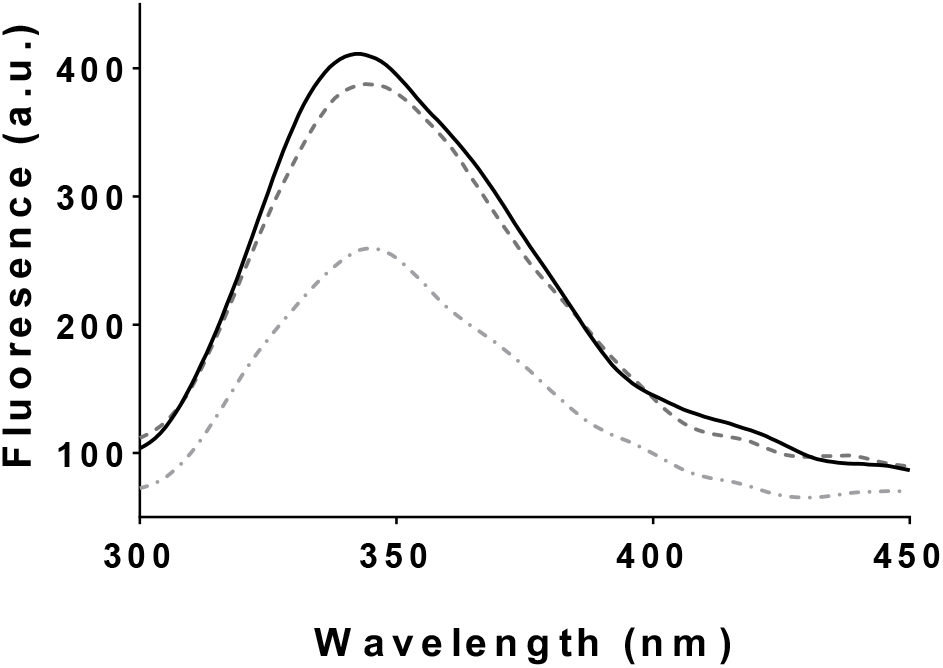
Variation of the fluorescence emission spectrum (λ_ex_=290nm) of the MBP317-347 switch enzyme in: absence of ligand (continuous line); presence of maltose (dashed line); or presence of alginate (dash-dotted line). Standard reaction conditions: 1 μM MBP317-347, 20 μM maltose or alginate, 30 mM phosphate buffer, pH 7.0, 20 °C.

Docking analysis was used to compare the binding of different alginate dimers in the maltose-binding pocket. These simulations predicted that the interaction between the maltose-binding pocket and alginate dimers is weaker than with the native ligand maltose, as shown in Table 1. This result would be expected, but the calculated binding energy for the alginate dimers (di-mannuronic and di-guluronic) was still energetically favorable. In contrast, the interactions with longer alginate oligomers are energetically unfavorable (Table 1). The amino acid residues that interact with maltose (Figure 3 a) are similar to those interacting with di-mannuronic acid (Figure 3b) only differing in the potential for hydrogen bond formation. In turn, the alginate hexamer must be folded completely to fit inside the maltose-binding pocket (Figure 3 c). The principal difference between MBP and the alginate binding protein (AlgQ2) is that the alginate-binding pocket of AlgQ2 is larger and deeper than the maltose-binding pocket of MBP.^12^ This means there is a high energy cost associated with folding the alginate chain inside the small maltose-binding pocket, in line with the unfavorable binding energy between alginate and MPB317-347.

**Figure 3.**
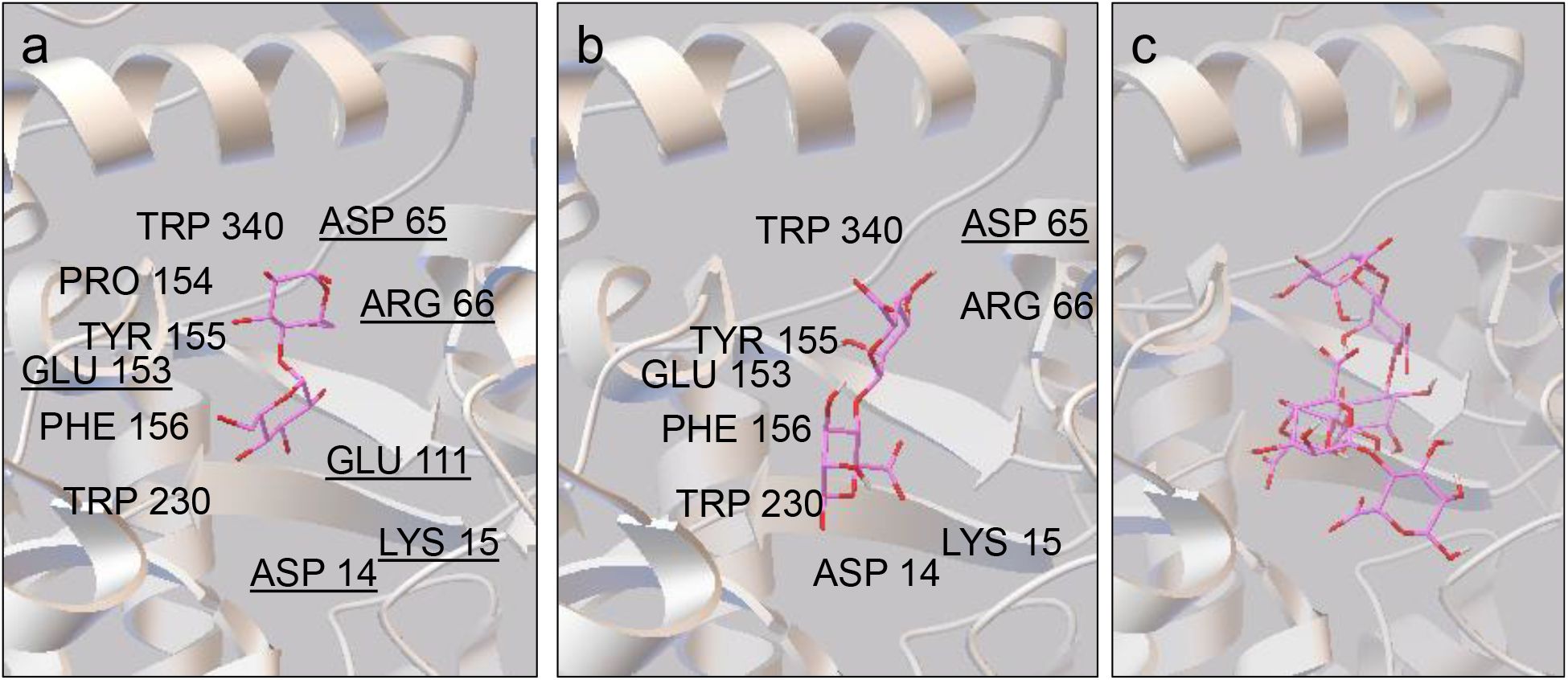
Molecular docking models of maltose-binding protein (PDB entry 1ANF) in complexes with different substrates, showing similar ligand coordinates. (a) Maltose; (b) Di-mannuronic acid; and (c) Mannuoric acid hexamer. Models generated using Autodock v4.2. Underlined amino acid residues are involved in hydrogen bonding between enzyme and substrate.

**Table 1.** Predicted binding energies (Autodock v4.2)

| Ligand | Binding Energy<sup>a</sup> | Binding Energy<sup>b</sup> |
| --- | --- | --- |
| Maltose | -6.8 | -7.5 |
| Mannuronic dimer | -1.4 | -3.2 |
| Guluronic dimer | -0.7 | -3.3 |
| Mannuronic hexamer | 2400 | 96.93 |

<sup>a</sup> Ligand position according to crystal structure (kcal/mol). <sup>b</sup> Optimal ligand position according to software (kcal/mol)

Different kinds of alginate-based nanoparticles were prepared to study their interaction with the switch enzyme. The particles were named according to their composition, namely, lysozyme-alginate (Lyz-Alg), lysozyme–alginate and lecithin (Lyz-Lec-Alg) and lysozyme–albumin–alginate and lecithin (Lyz/Alb-Lec-Alg). Table 2 shows the physical characteristics of the different formulations. The average diameters of the two formulations containing lecithin, with or without albumin, were similar (in the range 216–236 nm as determined by both DLS-NIBS and NTA) and they also shared a similar number of particles per unit volume. This is diagnostic that the physical dimensions and overall yield of formation of the nanoparticles is not affected by the presence of albumin. In turn, the lysozyme–alginate formulation (Lyz-Alg) was ∼100 nm larger in size but had a lower polydispersity index (PDI), and the values for number of particles were approximately half the corresponding values of nanoparticles containing lecithin (Table 2). The zeta potential (ζ) was highly negative in all three systems (∼ −45 mV), however nanoparticles comprising lecithin retained 20% more alginate.

**Table 2.** Physical characteristics of the different nanoparticle formulations at pH 4.5

| Formulation | DLS-NISB<sup>a</sup><br>(∅, nm) | PdI<sup>b</sup> | $\zeta$<br>(mV)<sup>c</sup> | Alg<br>(%) | NTA<sup>d</sup><br>(∅, nm) | [NPs]<sup>e</sup><br>(ps/ml) |
| --- | --- | --- | --- | --- | --- | --- |
| Lyz-Alg | 328±16 | 0.17 | -48±15 | 62±7 | 342±29 | 16±3 |
| Lyz-Lec-Alg | 216±8 | 0.24 | -44±4 | 87±5 | 223±9 | 36±8 |
| Lyz/Alb-Lec-Alg | 219±16 | 0.26 | -42±2 | 87±11 | 236±20 | 38±9 |

<sup>a</sup>Diameter by intensity as measured by dynamic light scattering with non-invasive back scattering (DLS-NIBS); <sup>b</sup>Polydispersity index; <sup>c</sup>Zeta potential values measured in water; <sup>d</sup>Diameter by number as measured by nanotracker analysis (NTA). <sup>e</sup>Nanoparticle count number per unit volume (value $\times 10^8$ particles/mL). Values are mean average ± standard deviation, ( $n = 3$ )

We investigated the interactions between the different nanoparticle formulations and the switch enzyme MBP317-347 to determine the suitability of the formulations as nanocarriers. In Figure 4, we show the activity of the enzyme in presence of the different nanoparticles in buffers with different ionic strengths (12 or 100 mM NaCl) and at pH values above and below pH 5.8, which is the calculated theoretical pI of MBP317-347. The increase in enzymatic activity observed in the presence of nanoparticles (Figure 4a) was an unexpected finding. Indeed, conformational changes of enzymes in presence of solid nanoparticles19 and or polyelectrolyte complexes.^20^ have been reported to cause a loss of activity. This has been attributed to partial unfolding by absorption on the surface or sequestration into a core that differs from the bulk solution.^21, 22^ Interestingly, adding NaCl to the system or increasing the pH only partially quenched the increase in enzyme activity caused by the nanoparticles (Figure 4b and 4c). Quenching was stronger at pH 7.0, where MBP317-347 should adopt a net negative charge, like the particles themselves. The increase in enzymatic activity may be explained by electrostatic interactions between the protein and particles bearing opposite partial charges, but such interactions are weak and do not persist in the presence of NaCl. Instead, the observed partial resistance of the enzymatic activity to addition of NaCl and pH suggests a more specific interaction between the alginate in the surface of the nanoparticle and the maltose-binding pocket of MBP317-347. The observed differences in activity between the three formulations may reflect subtle differences in the quantity or nature of alginate exposure on the particle surface, as also suggested by the enzymatic degradation experiment documented in our accompanying paper (ref).

**Figure 4.**
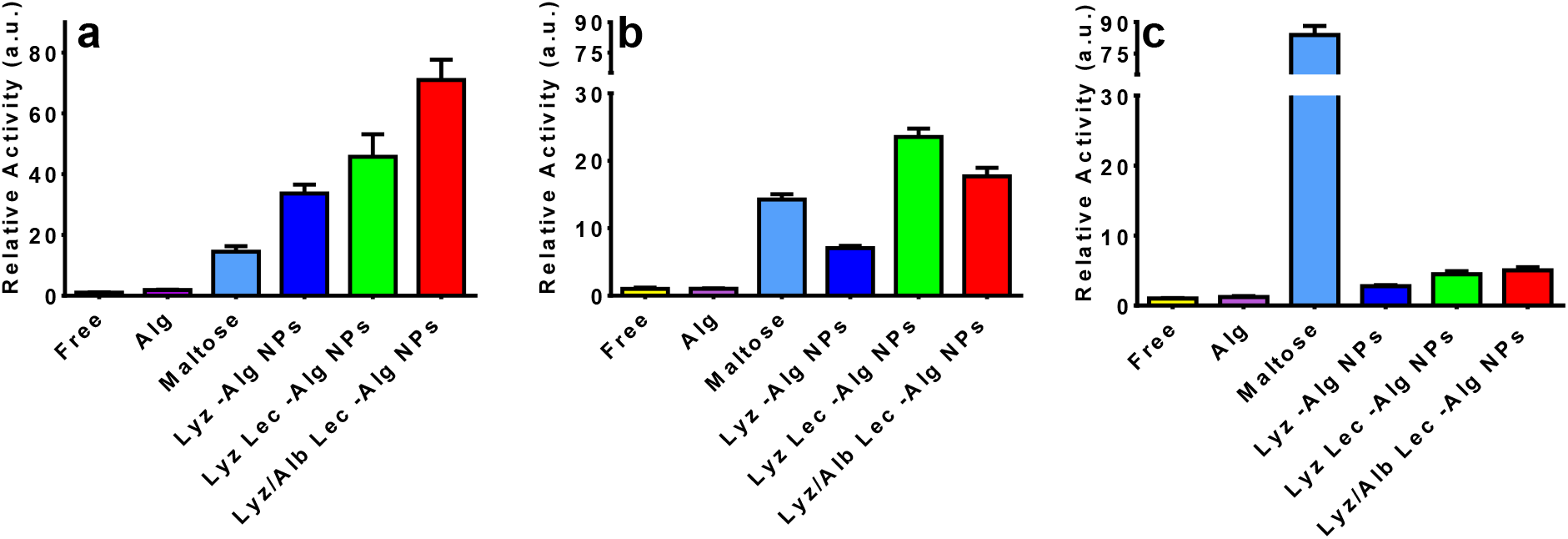
Relative enzymatic activity of MBP317-347 based on the hydrolysis of 50 μM nitrocefin in the absence (“free’) or presence of different allosteric ligands at the same disaccharide concentration (233 μM). (a) Reaction at pH 5.0. (b) Reaction at pH 5.0 with addded 100 mM NaCl. (c) Reaction at pH 7.0. Standard reaction conditions: 50.6 μM MBP317-347 in 30 mM phosphate buffer and 12 mM NaCl at 30°C.

Table 3 shows the affinity of the different effectors at pH 5.0 after analysis of the binding curves (Figure S1). The apparent *K_d_* of the enzyme for maltose is in the micromolar range whereas for alginate it is in the millimolar range, as previously observed for glucose.^10^ In contrast to the low affinity of alginate for the switch enzyme, the *K_d_* of it for the three alginate nanoparticle formulations is in the micromolar range, and the Lyz-Alg particles showed a binding constant that was similar to that of maltose. The large surface/volume ratio of the nanoparticles favors the exposure of an already immobilized high concentration of alginate molecules per unit volume, thus reinforcing the electrostatic attraction between the particle and the enzyme, and favoring stronger binding to the alginate. Since alginate is already confined at the nanoparticles, this may provide a thermodynamic advantage to the overall cost of loss of entropy involved in the formation of the enzyme-ligand complex. The lower affinity of the particles containing lecithin, despite the presence of more alginate on the surface (Table 1), suggests that the alginate is less readily accessible as an allosteric ligand. The maximum enzyme activity caused by each of the different effectors also correlated with the maximum fraction of enzyme bound (see kinetic parameters in Table 3) A different alginate distribution and conformation in the Lyz/Alb-Lec-Alg particles may explain the higher enzymatic activity observed in this system.

**Table 3.** Binding and kinetic parameters obtained after fitting the data from Figure S1 to the one-site-specific binding model (Standard reaction conditions: 50.6 nM MBP317-347, 30 mM phosphate buffer pH 5.0 and 12 mM NaCl, 20 °C).

| Formulation | $K_d^*$ (μM) | $V_{max}$ (μM/s) ** | $R^2$ |
| --- | --- | --- | --- |
| Lyz-Alg | 3.5±0.5 | 0.093±0.002 | 0.9802 |
| Lyz-Lec-Alg | 11.9±2.2 | 0.096±0.004 | 0.9869 |
| Lyz/Alb-Lec-Alg | 11.3±1.2 | 0.115±0.003 | 0.9599 |
| Alginate | 2409±269 | 0.010±0.010 | 0.9650 |
| Maltose | 2.9±0.3 | 0.043±0.001 | 0.9755 |

\*Apparent dissociation constant; \*\* Maximum activity.

The proportion of MBP317-347 associating with the particles was 49 ± 6% in the presence of maltose and 39 ± 8% in its absence, as determined by quantitative SDS-PAGE (Figure S2). This difference was not statistically significant (*p* = 0.14), thus confirming that maltose binding to the enzyme does not trigger its release from the particle surface. Is worth to mention that for thermodynamically controlled systems such as the one here described, it has been reported that ultracentrifugation produces stress in the formulations and destabilizes the nanosystems. Therefore, ultracentrifugation of our systems could have affected the interaction forces between the switch enzyme and the nanocomplex, thus leading to results that do not necessarily represent the condition in solution.^23^

Figure 5 shows the time-dependent activity of the enzyme in experiments carried out at 37 or 4°C in the presence of Lyz/Alb-Lec-Alg nanoparticles (Figure 6a), maltose (Figure 6b) and in absence of both (Figure 6c). The half-life of the activity of the enzyme at 37°C was 1.5-fold lower than at 4°C and more than the 80% of the activity is lost after 8 h at both temperatures. The presence of maltose increased the half-life of the enzyme from 5.2 ± 0.6 to 10.2 ± 2.7 at 4°C. In contrast, in the presence of the nanoparticle formulations, the enzyme activity remained constant for up to 8 h independently of the temperature, with only 8% of activity lost only after 31 h. The association between the protein and the alginate on the particle surface may therefore help to maintain the protein in its optimally-folded conformation. The interaction between the maltose-binding pocket and alginate may also protect the enzyme, as previously shown for maltose.^24^

**Figure 5.**
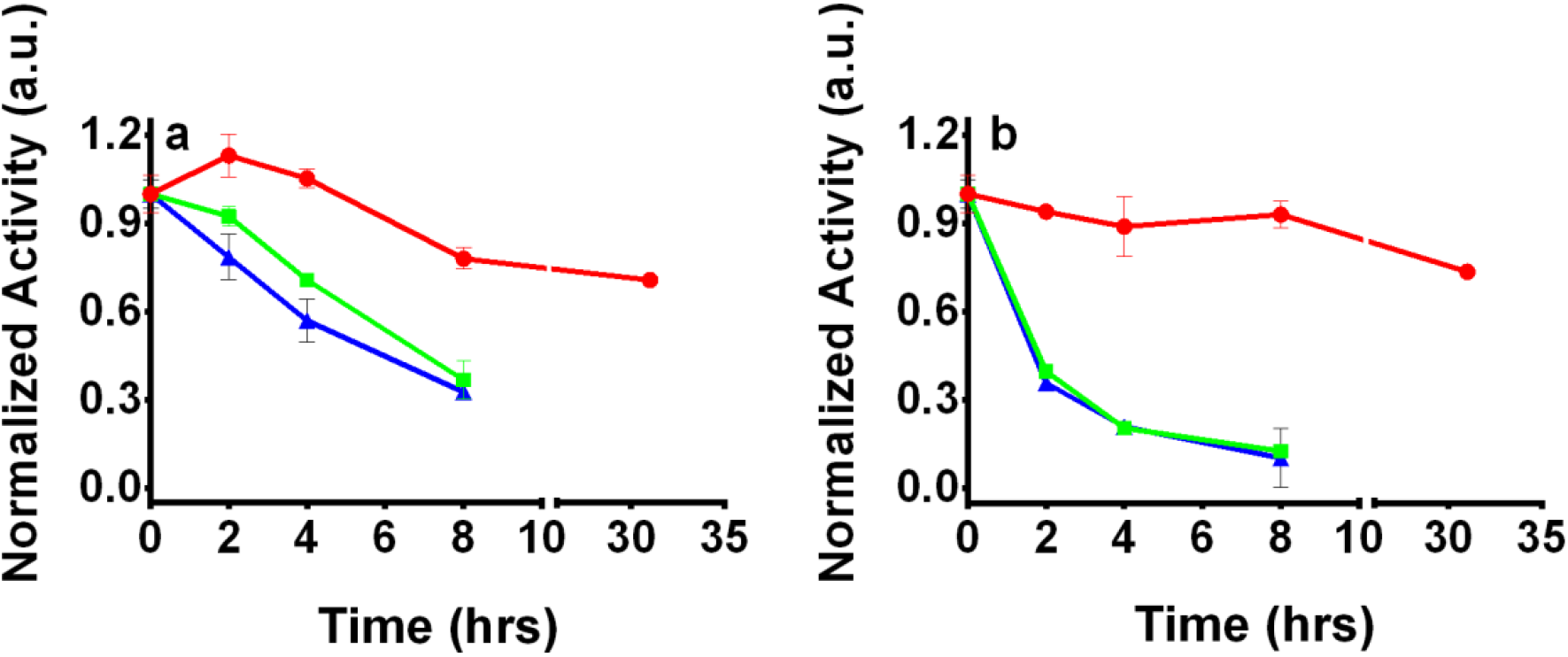
Enzymatic normalized activity of MBP317-347 at: a) 4 °C or b) 37°C, during incubation with:Lyz/Alb-Lec-Alg (red), maltose (green), or no ligand (blue) Standard reaction conditions: 50% PBS pH 7.3, 10% glycerol, 50.9 nM MBP317-347, maltose 233 μM and each formulation present at 233 μM disaccharide units.

## Conclusions

Alginate-protein particles with and without lecithin show the ability to increase the catalytic activity of MPB317-347 in different conditions. By contrast, alginate in solution had only a very minimal effect catalytic activity. The observed nanoparticle-dependent increase on enzymatic activity of the switch enzyme, the partial resistance of this increase to the addition of NaCl and an increase in pH, and the fluorescence quenching and docking studies support an important role for the maltose-binding pocket in the interaction between MBP317-347 and the alginate in all the studied nanoparticle systems. Finally, albumin-containing particles were found to improve the thermal stability of MPB317-347.

## Acknowledgements

We acknowledge support from the DFG, Germany (Project GRK 1549 International Research Training Group ‘Molecular and Cellular GlycoSciences’), The Danish Agency for Science, Technology and Innovation, Denmark (FENAMI project 10-093456) and the National Institute of General Medicine at the United States National Institutes of Health (R01 GM066972).

