## Supporting information for "Protein-surfactant-polysaccharide nanoparticles increase the catalytic activity of an engineered β-lactamase maltose-activated switch enzyme"

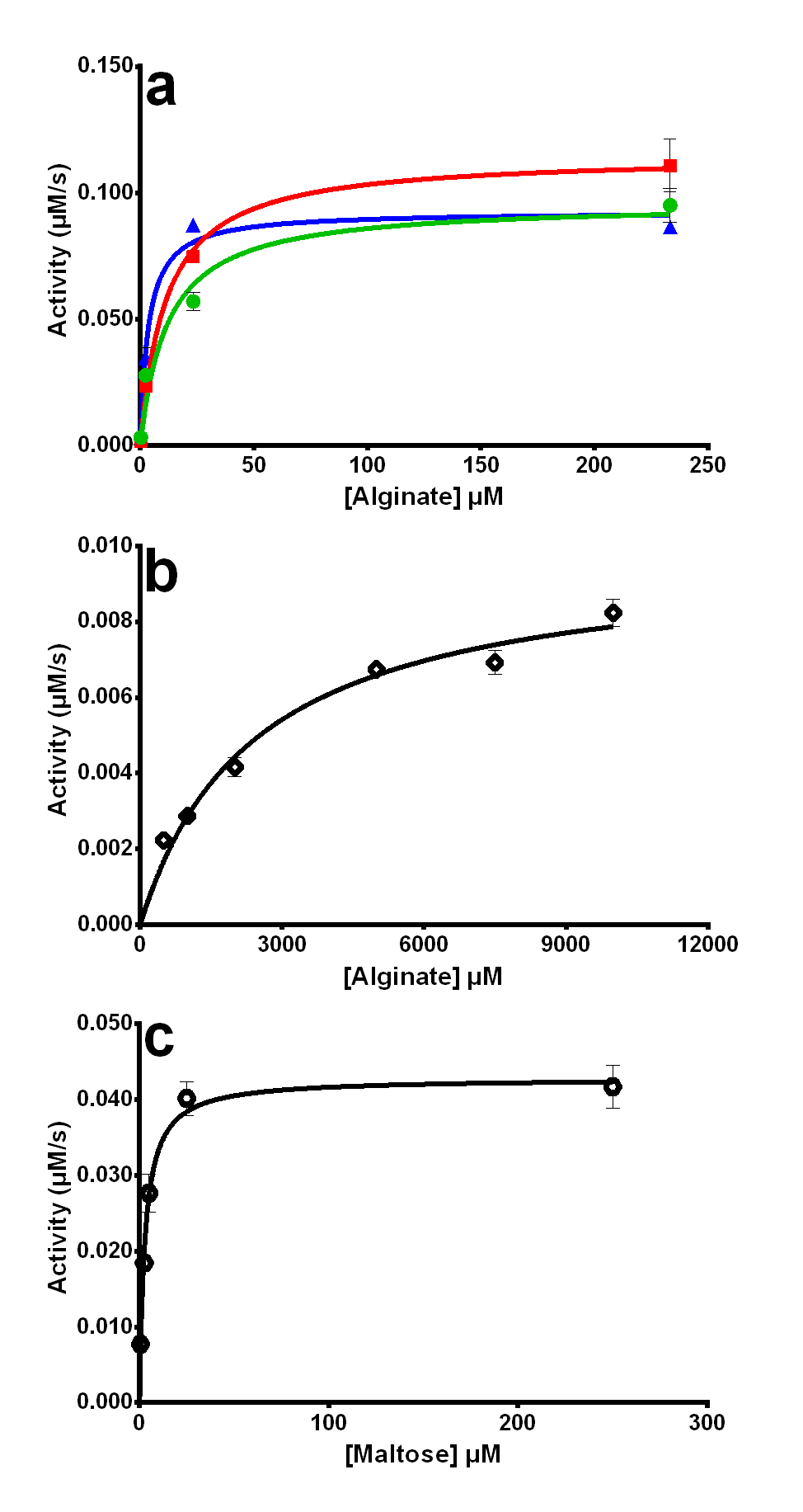

Figure S1. Enzymatic activity of MBP317-347 based on the hydrolysis of 50 µM nitrocefin in the presence of increasing concentrations of effector. (a) The three nanoparticles formulations (blue triangle) Lyz-Alg, (green dots) Lyz-Lec-Alg, (red square) and Lyz/AlbLec-Alg. (b) Alginate polymer in solution. (c) Maltose in solution. Standard reaction conditions: 50.6 µM MBP317-347 in 30 mM phosphate buffer and 12 mM NaCl at 30°C.

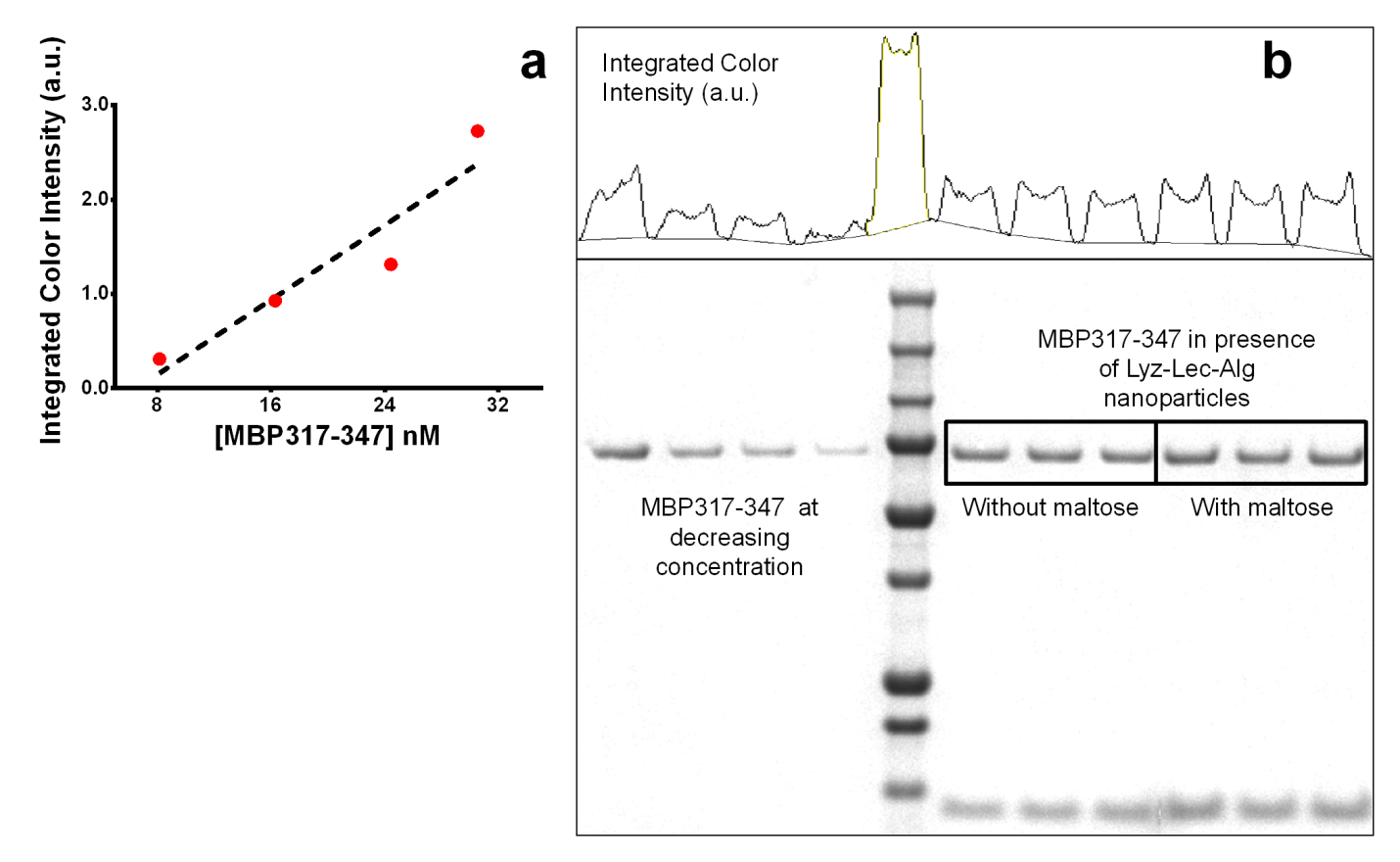

Figure S2. Gel retention assay: a) Example of a calibration curve from an ImageJ (National Institutes of Health) processed SDS-gel image of MBP317-347; b) Scanned image of SDS-gel of MBP317-347 at decreasing concentration and supernatant of MBP317-347 (50.9 nM) in presence of Lyz-Lec-Alg nanoparticles (disaccharide concentration 233 µM ) with and without maltose (100 µM ). The densitometric scan above the gel represents the color density within the bracketed region, which was further integrated with ImageJ (National Institutes of Health)
